# Reevaluation of the RNA binding properties of the *Tetrahymena thermophila* telomerase reverse transcriptase N-terminal domain

**DOI:** 10.1101/621573

**Authors:** Christina Palka, Aishwarya P. Deshpande, Michael D. Stone, Kathleen Collins

## Abstract

Telomerase restores chromosome-capping telomeric repeats lost with each round of genome replication by DNA-templated DNA polymerases. The telomerase reverse transcriptase (TERT) N-terminal (TEN) domain is a peripheral, telomerase-specific, processivity-stimulatory addition to more conserved domains that encircle the active site cavity. Reports of ciliate, yeast, and mammalian telomerase TEN domain associations with the telomerase RNA subunit (TR) describe low affinity interactions of uncertain specificity. Unfortunately two cryo-EM structures of synthesis-paused telomerase holoenzymes lack sufficient resolution to discriminate molecular specificity of possible TR contact(s) with the TEN domain, and there is no assigned density for the TEN domain termini implicated in RNA binding. Furthermore, studies have revealed alternative secondary structures for TR regions that could interact with TERT prior to TR folding into active conformation. Informed by recent advances in knowledge of telomerase structure, we returned to the investigation of *Tetrahymena thermophila* TERT TEN domain interaction with TR. Instead of finding specificity for a particular TR sequence or structure, we discovered that the tagged TEN domain used in previous characterizations has trace contamination with a bacterial RNA-interacting protein not detectable by SDS-PAGE. By resolving this interference, we show that the TEN domain binds RNAs dependent on RNA length rather than sequence.

## INTRODUCTION

Telomerase and other reverse transcriptases (RTs) discriminate their appropriate templates using sequence- and structure-specific RNA recognition. Retroelement RTs use their entire bound RNA as template [1]. In contrast, telomerase copies only a short region within the TR subunit of an active ribonucleoprotein (RNP) complex [2]. Telomerase biogenesis stably co-folds TERT and TR in a hierarchical series of induced conformational changes that ultimately determine the region of TR accessible to the active site [3]. Template 5’-boundary fidelity is essential for precise repeat synthesis and is strictly enforced in most but not all telomerase holoenzymes [4]. This boundary is set by hindrance from 5’ template-flanking RNA secondary structure or RNA-protein interaction [5, 6], and in vertebrate enzymes it depends in large part also on sequence-specific recognition of the template-product duplex [7, 8].

Critical RNA interactions of TERT and retroelement RTs are mediated by a protein domain immediately preceding the ubiquitously conserved RT motifs (Figure 1A). Within TERT, this high-affinity telomerase RNA binding domain (TRBD) binds and positions both a template 5’-flanking region (Stem II of ciliate TR, Figure 1B) and a distant, independently folded stem-loop motif (Stem IV of ciliate TR, Figure 1B) [6, 9–14]. TRBD interfaces with TR establish the tertiary structure of activity-essential TR motifs in the catalytic core of ciliate and human telomerase holoenzymes [14, 15]. Together the TERT TRBD, the RT domain with active-site motifs (RT), and the following TERT C-terminal extension (CTE) form the “TERT ring” encircling the active site cavity (Figure 1A)[16, 17]. Placement of the template in the vicinity of the active site cavity requires the template 3’-flanking region to traverse to the opposite side of the TERT ring from the TRBD-bound template 5’-flanking region, and for the TR path 3’ of the template to ultimately encircle the entire circumference of TERT ring (Figure 1C)[12, 14, 15]. The structural determinants of most of the TR path, including that of the template 3’ flanking region, are not yet established.

**Figure 1.**
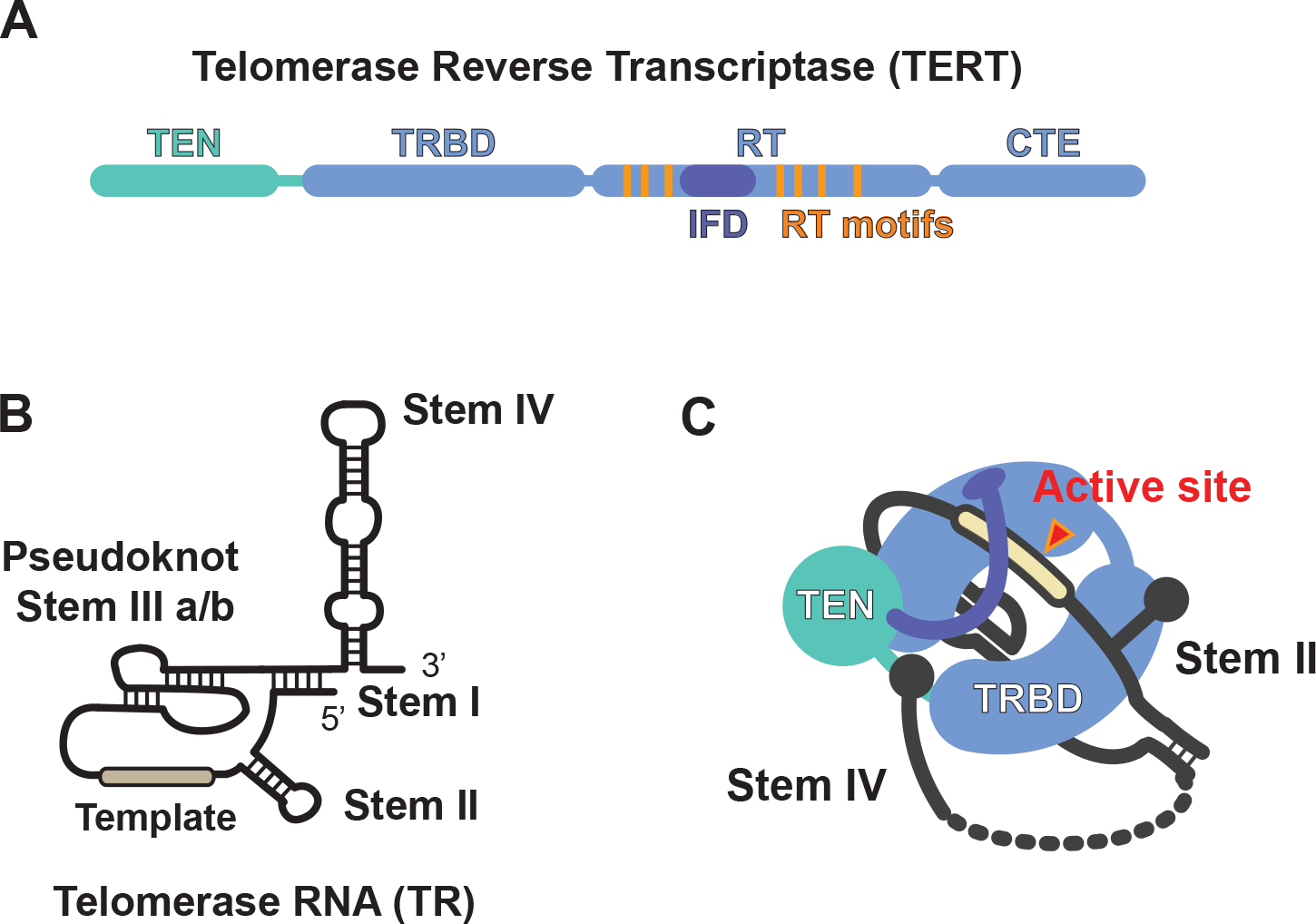
Conserved telomerase subunits from the ciliate *Tetrahymena thermophila*. (A) A schematic illustration of TERT domain organization, including: the Telomerase Essential N-terminal (TEN) domain, telomerase RNA binding domain (TRBD), the reverse transcriptase (RT) domain, and the telomerase C-terminal extension (CTE). The conserved insertion of fingers (IFD, purple) motif within the RT domain and canonical RT motifs (orange) are depicted. (B) Secondary structure of *Tetrahymena* TR with conserved stem elements and RNA pseudoknot. The region of TR that serves as the template during telomere repeat synthesis is demarcated in beige. (C) Cartoon model of the three-dimensional architecture of *Tetrahymena* telomerase RNP based on cryoEM structure [15, 30]. The model highlights the complex topological arrangement of the protein and RNA within the assembled RNP. Colors in model are as described in panel (A).

In telomerase holoenzymes of the ciliate *Tetrahymena thermophila* (henceforth *Tetrahymena)* and human cells, the TERT TEN domain is perched atop the TERT ring off to the RT-CTE side, instead of above the physically connected TRBD (Figure 1A and 1C)[14, 15]. The TR template 3’-flanking region threads past one side of the TEN domain as TR wraps around the CTE to the opposite face of TERT (Figure 1C). *Tetrahymena* and human TR take a generally similar path, despite complete divergence of the template 3’-flanking region sequence and structure. In *Tetrahymena* TR the template 3’-flanking region is entirely single-stranded, but in human TR the template is followed by a short single-stranded stretch and a long vertebrate-specific paired region [3, 8].

Ciliate, yeast, and human TEN domains can be expressed autonomously from the rest of TERT, and when purified they have been reported to interact with single-stranded DNA and/or TR as assayed by native gel electrophoresis, filter binding, cross-linking, NMR, and single-molecule assays [18–26]. Functionality of the recombinant, isolated TEN domain is supported by its co-assembly with TEN-less TERT and TR to reconstitute full enzyme activity [27, 28]. However, this complementation requires co-expression or co-assembly in a cell extract [28, 29], suggesting that conformational changes of the autonomously folded TEN domain may be necessary for productive protein-protein or protein-RNA domain interactions. Indeed, structures of the TERT TEN domains from *Tetrahymena* and the thermophilic yeast *Hansenula polymorpha* reveal disordered regions likely to be constrained within a fully assembled holoenzyme [18, 26].

The isolated *Tetrahymena* TEN domain was reported by two groups to bind *Tetrahymena* TR, with at least 100-fold lower affinity than the nanomolar binding of the *Tetrahymena* TERT TRBD [18, 21]. Initial studies from the Collins lab suggested that TEN domain interaction with TR had some dependence on TR sequence, but different non-overlapping TR truncations and internal deletions all strongly compromised binding, indicating that no single TR region sufficient for interaction [21]. Other studies suggested general RNA binding activity of the *Tetrahymena* TEN domain, which was shown to be dependent on the N- and C-terminal extensions of the folded TEN domain core that have not been resolved in any RNP structure to date [18, 30]. With recent improvements in knowledge about telomerase RNP architecture [14, 30], we sought to better characterize *Tetrahymena* TEN domain interaction with TR. We conclude that the isolated TEN domain has lower affinity for TR than previously reported and that it lacks obvious sequence specificity of interaction. Overall, TEN domain structure/function relationships remain an elusive goal for future understanding.

## RESULTS

### A bacterial contaminant with RNA binding activity co-purifies with 6xHis-tagged TEN domain

Previous studies bacterially expressed and purified an N-terminally six-histidine (6xHis) tagged *Tetrahymena* TERT TEN domain to investigate its TR and DNA binding activities, atomic resolution structure, and requirements for functional complementation with *Tetrahymena* TERT ring and other telomerase holoenzyme subunits [18, 21, 22, 28]. Here we purified this polypeptide using a similar process of affinity chromatography with nickel resin. TEN domain was assessed for purification from bacterial proteins by SDS-PAGE and Coomassie staining (Figure 2A, lane 1) and for RNA binding by electrophoretic gel mobility shift assays (EMSAs). In assays with limiting radiolabeled TR (~1 nM) and a large excess of 6xHis-TEN protein (micromolar range), the nickel-affinity (Ni-affinity) purified TEN domain appeared to bind TR with affinity slightly less than micromolar (Figure 2B), consistent with previously reported binding affinities [18, 21]. Our binding assays contained a vast excess of yeast tRNA intended to compete for any non-specific interactions (see Methods). In a separate experiment, we assessed the ability of unlabeled (cold) TR to compete for binding of radiolabeled TR. Unexpectedly, we observed a strong reduction in the amount of TEN-TR complex with sub-stoichiometric levels of cold competitor TR (Figure 2C), for example with only 75 nM cold TR added to 750 nM TEN domain. This result is unexpected because a higher concentration of cold TR should be necessary to saturate RNA binding to the TEN domain and thus exclude radiolabeled TR. One possible explanation for this observation was that the fraction of 6xHis-TEN competent to bind TR is very low, perhaps due to alternative protein conformation and/or aggregation. To address the state of protein aggregation, we further purified 6xHis-TEN by gel filtration chromatography. Contrary to our expectations, the well-defined peak of 6xHis-TEN at the monomer retention time of a Superdex 200 column exhibited less TR binding than the input when normalized to TEN domain concentration (data not shown). This raised the possibility that a contaminant protein other than 6xHis-TEN might be responsible for the observed mobility shift.

**Figure 2.**
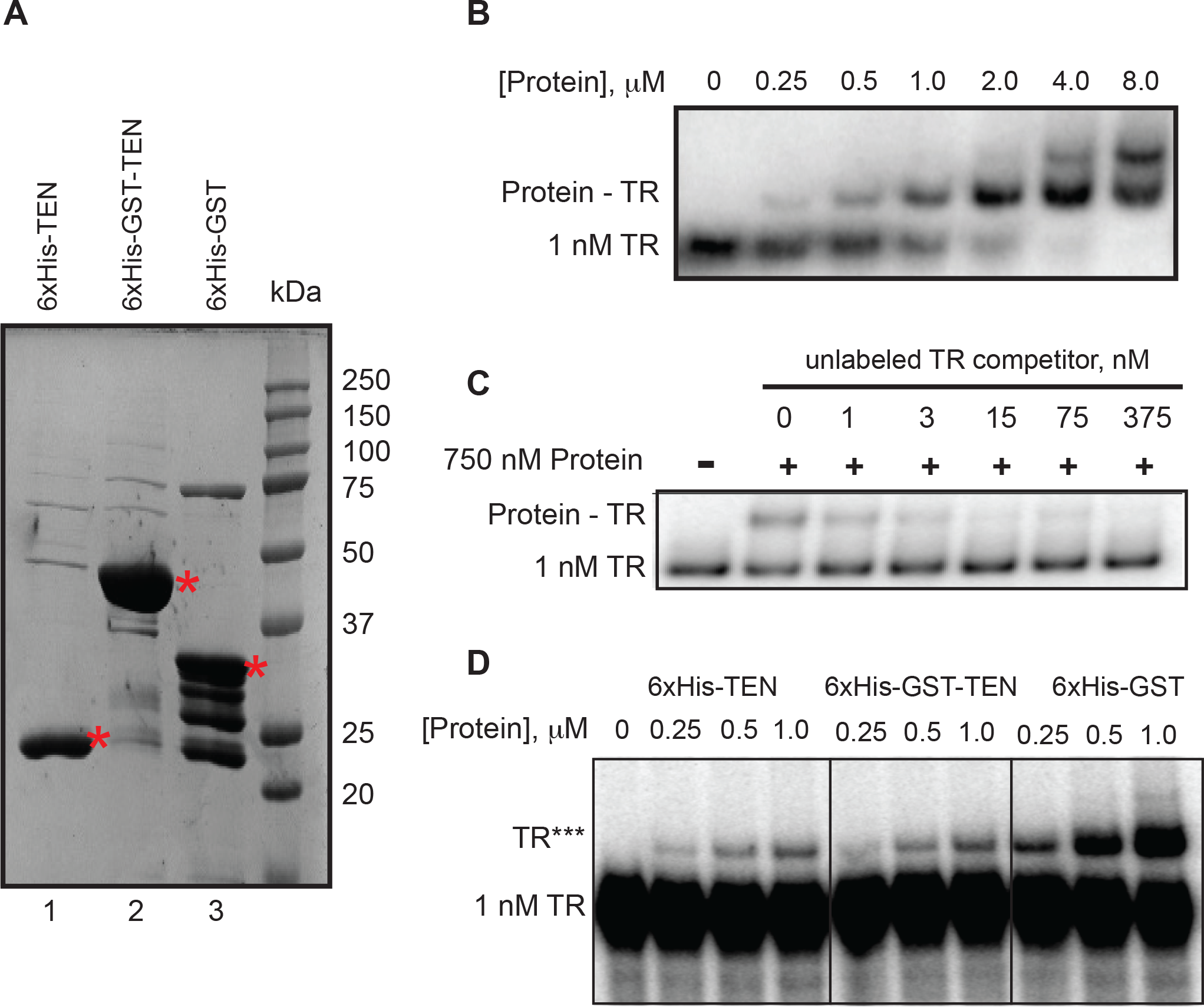
Purification and EMSAs testing GST fusion to the TEN domain. (A) SDS-PAGE analysis of 6xHis-TEN, 6xHis-GST-TEN, and 6xHis-GST (no TEN) proteins purified by binding to nickel resin. For each EMSA experiment, molar protein concentrations were calculated based upon the molecular weight of each fusion protein. Full length proteins are indicated by a red asterisk. (B) EMSA analysis of 1 nM radiolabeled TR incubated with the indicated amounts of 6xHis-TEN protein. (C) Competition EMSA experiment with 750 nM protein and 1 nM radiolabeled TR (~ 50% of TR bound). Unlabeled TR competitor was added to the binding reaction at the indicated concentrations. (D) EMSA of the indicated proteins with 1 nM radiolabeled TR. Proteins were normalized to the same molar concentration and added at the indicated concentrations. TR*** indicates a mobility shift of the radiolabeled TR.

To explore this possibility, we more than doubled the molecular weight of the TEN-domain polypeptide by inserting a glutathione S-transferase (GST) tag immediately following the 6xHis tag at the TEN domain N-terminus. We purified 6xHis-GST-TEN, as well as a 6xHis-GST negative control, using the same Ni-affinity method employed for 6xHis-TEN. SDS-PAGE analysis showed the expected sizes for the GST fusion proteins (Figure 2A, lanes 2-3). Protein samples were normalized to the same molar fusion-protein concentration and titrated into binding reactions with limiting TR (~1 nM). The purified protein samples all gave the same mobility shift (indicated as TR***), which was maximal in amount with the negative control 6xHis-GST sample (Figure 2D). This result implicates a bacterial contaminant rather than the *Tetrahymena* TEN domain as the source of TR mobility shift here, and by extension likely in previous RNA binding assays as well, even though no candidate for such a contaminant protein has been evident by SDS-PAGE in proportion to the mobility shift activity.

### Elevated TR concentration enables detection of RNA binding by the TEN domain

In additional experiments, we used different TEN domain fusion proteins and different assay conditions to investigate TR interaction with the *Tetrahymena* TEN domain. To this end, in parallel, we expressed and Ni-affinity purified 6xHis-TEN and TEN domain N-terminally tagged with 6xHis also bearing a maltose binding protein (MBP) tag at its C-terminus (6xHis-TEN-MBP) to nearly triple the mass of the fusion protein relative to 6xHis-TEN alone. Also in parallel we purified two negative control samples: TEN domain with C-terminal MBP tag but no 6xHis tag (TEN-MBP) and N-terminally 6xHis-tagged MBP (6xHis-MBP). All of the fusion polypeptides were soluble and all but the TEN-MBP fusion protein were enriched by Ni-affinity chromatography, as determined by SDS-PAGE and colloidal Coommassie staining (Figure 3A). To use the negative control TEN-MBP purification, the sample eluted from nickel resin was normalized to the 6xHis-MBP sample by equivalent eluted volume rather than protein amount. First, as for the experiments described above, we used limiting radiolabeled TR (1 nM) and excess protein (1 μM) (Figure 3B). As described above for 6xHis-GST-TEN, the 6xHis-TEN-MBP protein did not change the position of mobility shift observed with 6xHis-TEN sample (Figure 3B, lanes 1-3). Although TEN-MBP lacking the 6xHis tag was not enriched by Ni-affinity purification, this negative control sample gave the maximal mobility shift intensity (Figure 3B, lane 4). On the other hand, the mobility shift was only marginally detectable for the negative control 6xHis-MBP sample (Figure 3B, lane 5), consistent with the EMSA signal arising from an RNA-binding contaminant enriched by Ni-affinity resin in competition with 6xHis-tagged recombinant protein. These results parallel results described above in suggesting that the predominant TR mobility shift does not correspond to TR binding by the TEN domain. Also, again, no candidate for the contaminant bacterial protein that mediates the TR shift is evident among the proteins detected by SDS-PAGE and gel staining.

**Figure 3.**
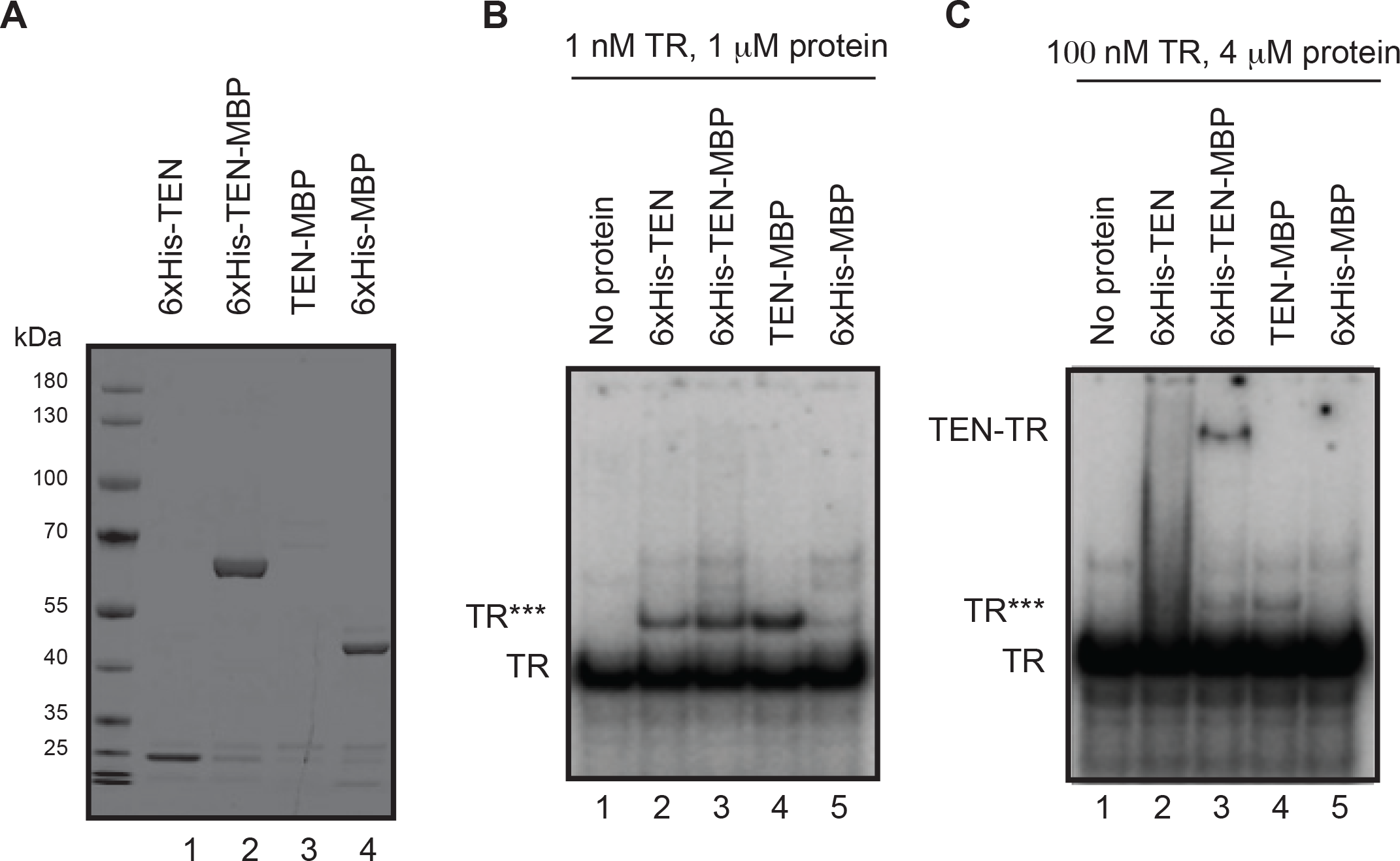
Purification and EMSAs testing MBP fusion to the TEN domain. (A) SDS-PAGE analysis of 6xHis-TEN (lane 1), 6xHis-TEN-MBP (lane 2), TEN-MBP (lane 3), and 6xHis-MBP (lane 4). (B) EMSA analysis of 1 nM radiolabeled TR with 1 μM of the indicated protein sample or control equivalent. TR*** indicates a mobility shift of the radiolabeled TR. (C) EMSA analysis of low specific activity 100 nM TR (99 nM cold TR plus 1 nM radiolabeled TR) with 4 μM of the indicated protein sample or control equivalent.

If the contaminant protein is of very low abundance, its TR mobility shift would disappear if limiting radiolabeled TR was mixed with unlabeled TR to generate lower specific activity RNA (as was indeed observed, Figure 2C). We therefore assayed the panel of purified proteins and controls for RNA binding using radiolabeled TR diluted with cold TR to a final TR concentration of 100 nM, mixed with 4 μM protein. In this binding condition, a different mobility shift was produced by 6xHis-TEN than by 6xHis-TEN-MBP, and neither of these shifts occurred with the negative control samples of 6xHis-MBP or TEN-MBP lacking a 6xHis tag (Figure 3C). The TR shift by TEN domain was more discrete for the protein with both 6xHis and MBP tags compared to the protein with 6xHis tag alone (Figure 3C, compare lanes 2 and 3), suggesting that MBP fusion improved TEN domain folding or its retention of RNA during gel electrophoresis. Taken together, these results demonstrate that lower specific activity TR at high concentration can be used to detect RNA binding by the TEN domain in EMSAs.

Having identified appropriate assay conditions to detect TR interaction with the TEN domain, we sought to characterize the interaction specificity. For this purpose we compared TEN domain binding to *Tetrahymena* TR with its binding to a streamlined version of human TR containing the activity-essential TR regions joined by a short linker (hTRmin, Figure 4A), which has a length about twice that of *Tetrahymena* TR (Figure 1B)[29]. Like *Tetrahymena* TR, hTRmin reconstitutes telomerase catalytic activity with its respective TERT translated in cell extract [29]. *Tetrahymena* and human TR primary sequences are largely divergent, and even shared secondary structure elements such as the pseudoknot vary in size (Figures 1B and 4A)[8]. In EMSAs, hTRmin was variably shifted in mobility by the panel of tagged *Tetrahymena* TEN domain proteins and negative controls. In assays with limiting 1 nM hTRmin and 1 μM TEN domain or control purification, the prominent mobility shift was the same in samples with 6xHis-TEN and 6xHis-TEN-MBP (Figure 4B, lanes 1-3). The negative control sample TEN-MBP gave maximal intensity of this mobility shift, and 6xHis-MBP gave the minimum, corresponding with results observed for mobility shift of *Tetrahymena* TR under conditions of limiting RNA (compare Figures 3B and 4B). In comparison, EMSAs using 100 nM lower specific activity hTRmin and 4 μM protein demonstrated mobility shift dependence on the TEN domain: a different mobility shift was produced by 6xHis-TEN than by 6xHis-TEN-MBP, and neither of these shifts occurred with the control samples (Figure 4C). These results parallel those obtained under the same binding conditions for *Tetrahymena* TR (Figure 3C). These findings suggest that the *Tetrahymena* TEN domain RNA-binding activity is not specific for *Tetrahymena* TR, concurring with a previous study [18], with the caveat that a limited number of RNAs have been tested. Although there is no evident primary sequence specificity, both the TEN domain and the bacterial contaminant appear selective for binding to RNAs at least half the length of *Tetrahymena* TR: a vast excess of tRNA did not compete binding (here and in previous EMSAs [21]), and subregions of *Tetrahymena* or human TR also did not compete (data not shown) [21].

**Figure 4.**
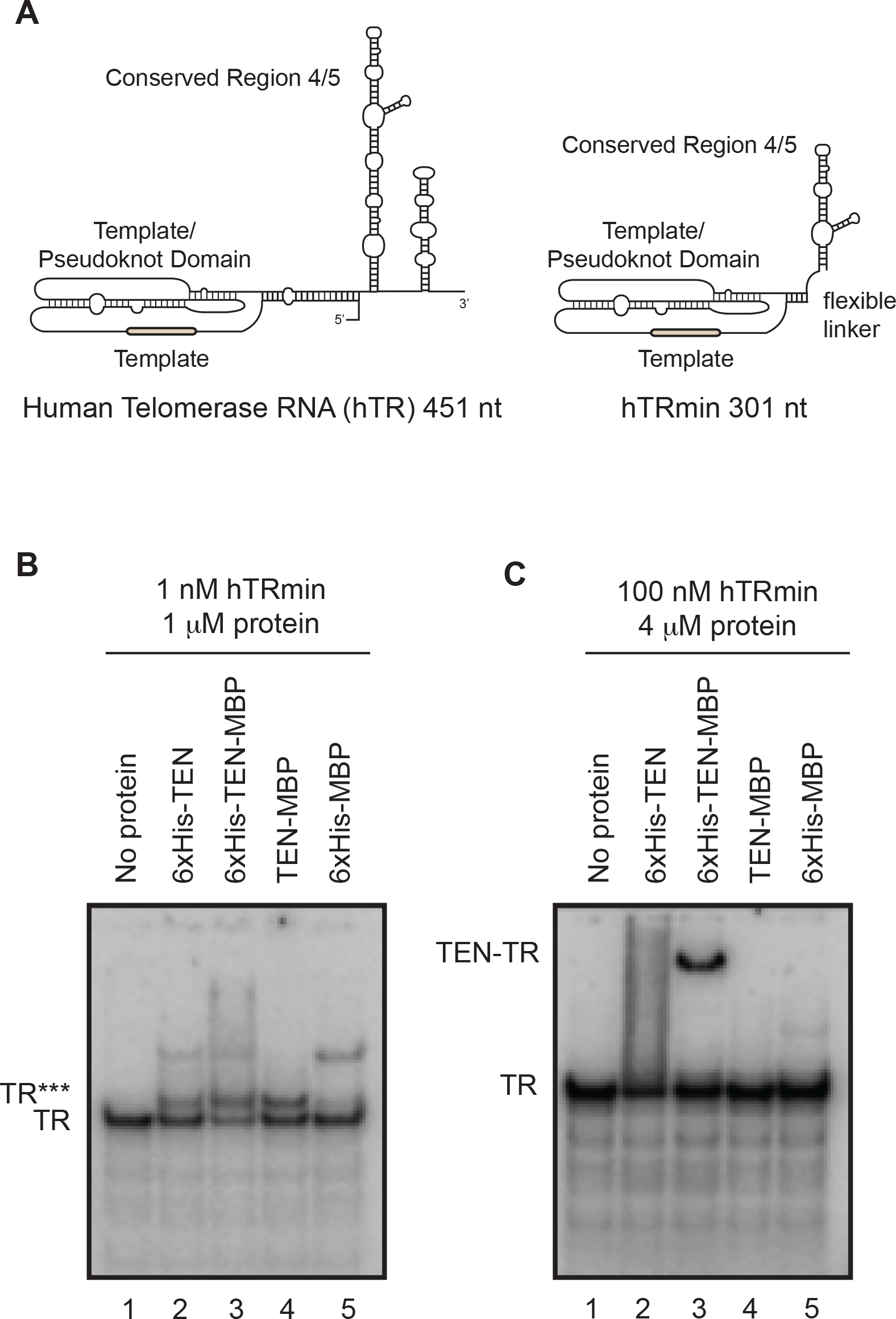
Comparison of RNAs for binding to the TEN domain. (A) Secondary structure schematics for full-length human TR (hTR) and a minimal activity-reconstituting human TR (hTRmin). The two activity-essential hTR regions are the Template/Pseudoknot domain and Conserved Region 4/5 [4, 8]. (B) EMSA analysis of 1 nM radiolabeled hTRmin with 1 μM of the indicated protein sample or control equivalent. TR*** indicates a mobility shift of the radiolabeled hTRmin. A low-abundance non-specific complex higher in the gel compared to TR*** is present in lanes with any 6xHis-tagged protein (lanes 2, 3, 5), suggestive of the presence of an RNA-binding contaminant that co-purified with any 6xHis-tagged protein. (C) EMSA analysis of low specific activity 100 nM hTRmin with 4 μM of the indicated protein sample or control equivalent.

## DISCUSSION

We show here that a trace amount of a bacterial protein with high affinity for TR can co-purify with 6xHis-tagged *Tetrahymena* TEN domain. Comparing across the purification samples described above and others, we did not find a protein evident by staining after SDS-PAGE that correlated in abundance with the amount of 1 nM TR mobility shift. A very low level of the contaminant is consistent with detection of a mobility shift only with high specific activity TR, for which we estimate ~40 pg of a 20 kDa protein contaminant would be sufficient, an amount less than 0.1% of the TEN domain. Mobility shift assays with lower specific activity TR were useful in detecting an RNA interaction authentic to the TEN domain. Although the trace-level contaminant still binds TR, its RNP becomes less abundant than the TEN domain RNP of interest.

Likely contamination of *Tetrahymena* TEN domain in previously published assays [21] prompts a revision of prior conclusions about TEN domain affinity and specificity of RNA binding. Similar to the RNA binding specificity of the bacterial contaminant, TEN domain interaction with TR appears sensitive to RNA length, perhaps due to RNA length-dependent formation of secondary structure. It is important to note that both the *Tetrahymena* and yeast TEN domain with determined structure possess functionally critical regions that are disordered [18, 26]. This could reflect the absence of interactions made in holoenzyme context: recent studies in several organisms have led to increasing appreciation of the TEN domain as a nexus of interactions between the catalytically active TERT ring RNP and telomerase holoenzyme and telomere proteins required for telomerase function and coordination at chromosome ends [31]. Because TEN domain conformation in active RNP may depend on the interacting proteins, properties of the autonomous TEN domain have uncertain significance for its roles in holoenzyme context.

Cryo-EM studies of *Tetrahymena* and human telomerase holoenzyme subunit architecture place the TEN domain above the TERT CTE, close to where the template 3’ flanking region begins its circumnavigation to the back side of TERT [12, 14, 30]. The *Tetrahymena* TR template 3’-flanking region changes dramatically in structure during TR folding with TERT [32–35], and likely during the catalytic cycle as well [31, 36]. For *Tetrahymena* TR, the mature RNA fold is not adopted without TERT: the template 3’ end, template 3’-flanking region, and some of the pseudoknot sequence form a long, snap-back hairpin [32–34]. Whether or not the TEN domain plays a role in the required refolding of TR to its active conformation remains to be addressed.

## METHODS

### Expression and purification of RNAs

*Tetrahymena* TR and human hTRmin were transcribed from linearized plasmids largely as previously described [29, 37], using T7 RNA polymerase, and then RNAs were purified by denaturing PAGE. RNA purity was verified by denaturing PAGE with SYBR Gold or ethidium bromide staining. RNA concentrations were determined by Nanodrop spectrometer (ThermoFisher).

### Expression and purification of proteins

*Tetrahymena* TERT TEN domain (amino acids 1-195) was expressed in fusion with a polypeptide tag or tag combination as indicated in the text, in parallel with expression of the large tags alone. All polypeptides were expressed using pET28 vectors in *E. coli* BL21(DE3) cells. Transformed cells were grown at 37°C until an optical density of approximately 0.6 was reached, at which point cultures were shifted to lower temperature for induction of protein expression by addition of approximately 1.0 mM isopropyl 1-thio-β-D-galactopyranoside. Aliquots of purified protein were stored at −80°C after flash freezing in liquid nitrogen.

For the experiments in Figure 2, protein was expressed by overnight induction at 18°C. Harvested cells were resuspended in TENA (20 mM Tris-HCl, 250 mM NaCl, 10 mM imidazole, 10% glycerol, 2 mM 1,4-dithiothreitol (DTT); pH 8.0). Cells were lysed via cell disruptor, after which slurry was clarified by centrifugation. Supernatant was mixed with Ni Sepharose Excel resin (GE Healthcare) pre-equilibrated with the lysis buffer and allowed to rotate end-over-end at 4°C for 3 hours. Resin was collected and washed at 4°C with approximately 10 column volumes of lysis buffer until no protein came off the column as determined by Bradford assay. Bound protein was eluted in 1 mL fractions with TENA adjusted to 250 mM imidazole. Protein was then dialyzed back into TENA lacking imidazole and concentration was determined by Nanodrop.

For the experiments in Figures 3 and 4, protein was expressed by 4 hour induction at room temperature. Harvested cells were washed with 1x PBS containing 200 μM phenylmethylsulfonyl-fluoride (PMSF) before freezing at −80°C. Thawed cells were resuspended in TENB (20 mM Tris-HCl, 2 mM MgCl_2_, 50 mM NaCl, 20 mM imidazole, 10% glycerol, 0.05% NP-40, 1 mM DTT; pH 8.0) with 200 μM PMSF, 1:200 of protease inhibitor cocktail (Sigma) and 1 mg/ml lysozyme. The slurry was gently stirred at 4°C for 45 min, followed by sonication for 3 min with 10-sec on/off pulses. Lysate was clarified by centrifugation, mixed with nickel-nitrilotriacetic acid-agarose resin (NiNTA, Qiagen) and allowed to rotate end-over-end at 4°C for 4 hours. Resin was collected and washed at 4°C with 3 changes of a large bead-volume excess of TENB for 15 min each. Bound protein was eluted in TENB adjusted to 300 mM imidazole. Protein concentration was determined by Bradford assay.

### EMSAs

For radiolabeling, purified RNAs were treated with shrimp alkaline phosphatase at 37°C for 1 h then end-labeled using T4 polynucleotide kinase and γ-^32^P-ATP at 37°C for 1 h. Complexes were resolved by electrophoresis at 4°C on native 5% acrylamide gels (37.5:1 acrylamide:bis acrylamide, 0.5x Tris borate-EDTA, and 4% glycerol added for gels in Figures 3 and 4). Gels were dried and exposed to phosphorimager screens. Products were visualized by scanning on a Typhoon (GE healthcare).

For the experiments in Figure 2, the binding reaction used binding buffer A unless otherwise specified (20 mM Tris-Base, 1 mM MgCl_2_, 50 mM NaCl, 10% glycerol, 1 mM DTT; pH 8.0) with 0.1 mg/mL yeast tRNA (Sigma) and 0.1 mg/mL BSA (NEB). Reactions were incubated on ice for 10 min. For the experiments in Figures 3 and 4, radiolabeled RNA was spiked into unlabeled RNA, heated to 70°C for 3 min, and slow-cooled to room temperature. RNA and protein were diluted in binding buffer B (20 mM Tris-HCl, 1 mM MgCl_2_, 100 mM NaCl, 10% glycerol, 5 mM DTT, 0.25 μl RNasin (Promega), trace bromophenol blue; pH 8.0) and incubated at room temperature for 20 min. Similar results were obtained with or without the presence of 0.1 mg/mL yeast tRNA (Sigma) and 0.1 mg/mL BSA (acetylated BSA from New England Biolabs).

